# Magnetoencephalographic Signatures of Conscious Processing Before Birth

**DOI:** 10.1101/2020.10.27.357129

**Authors:** Julia Moser, Franziska Schleger, Magdalene Weiss, Katrin Sippel, Lorenzo Semeia, Hubert Preissl

## Abstract

The concept of fetal consciousness is a widely discussed topic. In this study, we applied a hierarchical rule learning paradigm to investigate the possibility of fetal conscious processing during the last trimester of pregnancy. We used fetal magnetoencephalography, to assess fetal brain activity in 56 healthy fetuses between gestational week 25 and 40, during an auditory oddball paradigm containing first- and second-order regularities. The comparison of fetal brain responses towards standard and deviant tones revealed that the investigated fetuses show signs of hierarchical rule learning, and thus the formation of a memory trace for the second-order regularity. This ability develops over the course of the last trimester of gestation, in accordance with processes in physiological brain development and was only reliably present in fetuses older than week 35 of gestation. Analysis of fetal autonomic nervous system activity replicates findings in newborns, showing importance of activity state for cognitive processes. On the whole, our results support the assumption that fetuses in the last weeks of gestation are capable of consciously processing stimuli that reach them from outside the womb.

## 1. Introduction

Can a fetus be conscious? Not only has this question remained unsolved for many decades, it has also fostered numerous discussions (Anand et al., 1999; Lagercrantz and Changeux, 2009; Padilla and Lagercrantz, 2020). The answer to the question of fetal consciousness mainly depends on two aspects: First, how do we define consciousness and what are its neurophysiological correlates? Second, is conscious perception at the fetal stage possible in the light of fetal brain anatomy and physiology?

The concept of consciousness has many definitions. While the Oxford Advanced Learner’s Dictionary (2020) defines it as “the state of being able to use your senses and mental powers to understand what is happening”, researchers have proposed countless other definitions in the past decades. These range from consciousness as a neurobiological process to self-awareness and free will (for a summary see Vimal, 2009). In the last few years, research has challenged the borders of this anyhow vague concept even further by discussing self-consciousness of fish (Kohda et al., 2019) and the possibility of functional brains ex cranio (Vrselja et al., 2019). In light of these findings and the variety of possible definitions, it would seem that consciousness is not an all-or-nothing phenomenon but rather that it exists in different degrees. One such example of a more fine-grained concept is presented by Edelman (2003), who divides consciousness into primary and higher-order consciousness. Whereas primary consciousness mainly focuses on the integration of different sensory inputs together with memory, enabling one to adapt dynamically to a situation. For such a fine-grained concept of consciousness, the temporal scale on which an agent operates plays an important role. This has been developed further by Kent et al. (2019), who place different concepts of consciousness on a logarithmic temporal scale – from conscious experience in the range of seconds to self-consciousness in the range of months. In the context of this finely differentiated view, it seems plausible that some more primary forms of consciousness emerge as early as during the fetal period.

According to Kent et al. (2019), consciousness is tightly linked to memory, particularly to the ability to form memory traces that span a certain period of time. This is necessary to make predictions about events that take place in different temporal resolutions. Marchi and Hohwy (2020) relate the formation of predictions in the adequate temporal resolution to conscious processing. They pointed out that “a conscious agent is conscious of the hypotheses that are flagged at the appropriate resolution”, i.e. a conscious agent is able to adapt its predictions to the time-scale relevant for the current context.

With regard to this definition, it must first be clarified whether the fetal brain is generally capable of performing these processes. One key element for enabling conscious perception is the establishment of thalamo-cortical pathways in the cortical plate. During the midfetal period (15-23 weeks post conception) thalamo-cortical axons enter the intermediate zone and gradually accumulate in the subplate before moving to their cortical target regions (Kostović et al., 2019). The thalamo-cortical pathways are then established after week 24 of gestation and continue to grow in the late fetal period (early preterm phase 24-28 weeks), making cortical responses towards sensory input possible (Kostović et al., 2019; Kostović and Judaš, 2010; Lagercrantz, 2014). The latter divided the remaining gestational weeks after this key event into the late fetal period/late preterm phase (week 29-34) characterized by an intensive development of long associative pathways and the neonatal phase (commencing from week 35 post conception), during which the short cortico-cortical pathways essential for cortical functional connectivity are increasingly formed (Kostović et al., 2019).

Generally speaking, the number of long range connections increases with fetal age, and connectivity in brain networks is enhanced (Thomason et al., 2015). From gestational week 35 onwards, connections between regions belonging to the Default Mode Network (DMN) can be observed (Thomason et al., 2015). On the whole, the modularity of brain organization decreases with increasing gestational age (GA; Thomason et al., 2014).

This anatomical development over the fetal period indicates that neural networks necessary for conscious processing have been established before birth, which makes it prudent to investigate these processes during the last trimester of gestation.

Bekinschtein et al. (2009), who worked with patients with disorders of consciousness, introduced an auditory oddball paradigm for the investigation of memory traces over different time scales, the so called “local-global” paradigm. The local-global paradigm consists of sequences of tones which can either contain only identical tones or end with a different tone. At the same time, the sequence itself can be either frequent or rare. With this structure, the paradigm includes first order (within a sequence) and second order (across sequences) rule violations and therefore requires the formation of predictions on two hierarchical levels, operating on two time-scales. The neuronal correlate of the detection of an error in the prediction of the second-order rule can be measured by the event-related P300 component in the electroencephalogram. In healthy participants, the occurrence of a P300-like component showed the ability to detect a second order rule violation. Participants in an unresponsive wakefulness state however, were not able to show this component. The authors concluded that the ability to form a memory trace that is long enough to incorporate the second-order regularity – which goes hand-in-hand with the ability to form a prediction in the appropriate temporal scale – is a sign for conscious processing (Bekinschtein et al., 2009).

Apart from populations with disorders of consciousness, Basirat and colleagues (2014) showed that it is feasible to use this paradigm in three-month-old children. In fact, children were able to learn the hierarchical rules, as can be observed in the neuronal responses following the adequate temporal resolution (Basirat et al., 2014). In Moser et al. (2020), we successfully reproduced the same effect in newborns during their first eight weeks of life. The current paper therefore aims to take up the challenge to examine hierarchical rule-learning abilities in late gestation.

The implementation of auditory stimulation paradigms in fetal populations recorded with fetal magnetoencephalography (fMEG) has often shown fetal ability to respond to sounds from the environment (e.g. Eswaran et al., 2002; Schleussner et al., 2004; Wakai et al., 1996). In simple oddball paradigms, their ability to differentiate between two tones – and therefore to form simple predictions – was demonstrated (Draganova et al., 2005; 2007). Other, more complex paradigms revealed the fetal ability to habituate to a stimulus (Muenssinger et al., 2013) and to differentiate between different set sizes (Schleger et al., 2014).

The paradigms used in these studies enabled us to demonstrate the existence of basic cognitive abilities before birth. However, all these tasks require only a very short memory span (the preceding few tones) and do not take multiple temporal scales into account. It remains unclear as to whether the fetus is able to form a memory trace for second-order regularity in the last trimester of pregnancy and whether it adapts its predictions to the appropriate temporal scale. If a response towards second-order rule violations is observed, it implies that the fetus can integrate information over a longer time span and adapts its predictions on the basis of the rules learned. On the basis of anatomical development, this should be theoretically possible and, in the light of the aforementioned definition of consciousness, would be a sign of conscious processing.

To address this open question, the present study uses an auditory local-global oddball paradigm in fetuses between weeks 25 and 40 of gestation. By using such a large range of gestational ages, we wish to address two main questions: First, we wish to determine whether the ability of second-order rule learning is generally present during fetal life. Second, if it is present, we hope to determine whether it is present throughout the last trimester of pregnancy or whether it emerges at a certain point during development. In addition, we aim to explore the impact of fetal behavioral states on second-order rule learning by trying to replicate our findings from newborns whereby newborns in a more active state – as shown by a high heart rate variability (HRV) – had higher rule-learning abilities (Moser et al., 2020).

## 2. Material and Methods

### 2.1. Study Population

Participants were 60 mothers-to-be whose fetuses ranged from week 25 to 40 GA. Participants were recruited from Tübingen and the surrounding area. All mothers had uncomplicated singleton pregnancies and did not report any smoking or drinking. Four participants were not taken into consideration because their measurements were interrupted at an early stage, resulting in a study population of N=56 (GA range 25-40, mean = 32.39 ± 4.15 weeks, 25 female and 31 male fetuses). The present study combines cross-sectional and longitudinal recordings whereas 22 of the 56 participants were recorded multiple times (7 two times, 3 three times and 12 four times) in different gestational weeks resulting in 105 recordings in total. The local ethics committee of the Medical Faculty of the University of Tübingen approved the study and the consent form was signed by all participants. They received 10 Euro per hour for their participation.

Following data preprocessing, fetal brain activity could be detected in 81 recordings. These recordings included N=43 fetuses (GA range 25-40, mean = 32.96 ± 4.08 weeks; 19 were female, 24 male). Twenty-four fetuses were measured once and 19 had several measurements with four (N=5), three (N=9) or two (N=5) recordings (supplementary Table 1).

### 2.2. Material and Design

The material and design of this study were equivalent to those of Moser et al. (2020) and are also described here for the sake of completeness: stimuli were two pure tones (500Hz and 750Hz, duration = 200ms), presented at 90dB sound pressure level (approximately 60dB reaching the fetus after attenuation by the maternal tissue (Querleu et al., 1988)). One tone was assigned to be the standard (s) and one was assigned to be the deviant (d). Tones were presented in sequences of four, separated by a 400ms inter-tone-interval (total duration of the sequence = 2000ms). Each sequence was separated from the next by a 1700ms inter-stimulus-interval. Standard trials (ssss) consisted in the repetition of the same tone (i.e. the standard tone), whereas the other tone was introduced in the final position for deviant trials (sssd). Global rule violations are depicted by the upper case letter S or D (Figure 1).

**Figure 1:**
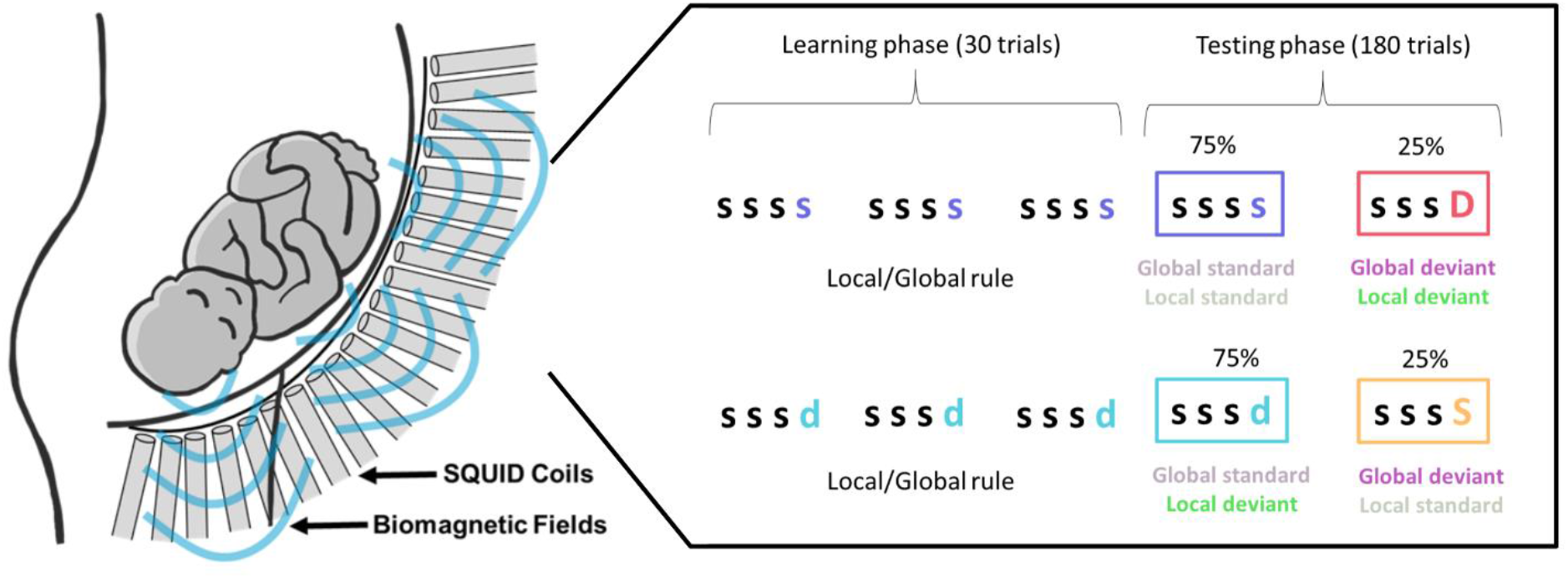
Left: schematic picture of fMEG recording. Right: experimental Paradigm. Each row represents one experimental block. Colors depict individual stimulus conditions. Right column describes the role of the sequence in the paradigm. Figure adapted from Moser et al. (2020).

Each block commenced with a learning phase during which a specific sequence (ssss or sssd) was repeated 30 times. This sequence was also the most frequent type of trials (75% occurrence) in the subsequent testing phase, the other type of sequence being randomly presented in 25% of trials. Two types of blocks were therefore possible: either 75% of the sequences were standard sequences (e.g. ssss ssss ssss) with 25% deviant sequences (sssD); or 75% of the sequences were deviant sequences (e.g. sssd sssd sssd) with 25% standard sequences (sssS). Two levels of mismatches were therefore assessed in this paradigm, at either the sequence (i.e. local) level (ssss vs. sssd) or the block (i.e. global) level (global standard, respecting the block rule (frequent ssss or sssd) vs. global deviant (rare sssS or sssD).

Each participant was recorded with the two types of blocks (creating two datasets per recording), and the order of blocks was counterbalanced across participants. At the beginning of each block, the learning phase implemented the global rule, followed by a testing phase with 180 trials comprising 25% global deviant trials (Figure 1). The trials in the testing phase were pseudorandomized, with a minimum number of two standard trials between deviant trials. This resulted in an overall duration of about 13 minutes per block. Both tones (500Hz and 750Hz) were used as standard tone or deviant tone across participants to control for a possible stimulus effect. In the event of longitudinal participation, the standard tone within one participant remained unchanged.

### 2.3. Experimental Procedure and Recording

#### Fetal magnetoencephalography

Fetal brain activity was measured using the SARA (SQUID Array for Reproductive Assessment, VSM MedTech Ltd., Port Coquitlam, Canada) fMEG system installed at the fMEG Center at the University of Tübingen. This is a non-invasive tool for measuring heart and brain activity of fetuses in the last trimester of pregnancy. The system, which contains 156 primary magnetic sensors, and 29 reference sensors distributed over a concave array designed to match the maternal abdomen, is installed in a magnetically shielded room (Vakuumschmelze, Hanau, Germany) to attenuate magnetic activity from the environment.

A sound balloon (King Systems Corporation, Noblesville, USA) can be placed between the maternal abdomen and the sensor array for the presentation of auditory stimuli. Fetal head position is determined prior to the measurement by ultrasound (Ultrasound Logiq 500MD, GE, UK) and marked by a localization coil placed on the maternal abdomen. To track maternal position changes in relation to the sensor array, three additional localization coils are placed on the spine, left and right side of the participant. Fetal head position is again determined by an ultrasound directly after the recording. In the event of a major change in position, the datasets are not taken into consideration.

#### Procedure

A timeslot of two hours was allocated to each participant. Upon arrival at the lab, an anamnesis was performed by our in-house midwife. All participants were informed about the aims of the study and signed the participant consent form. Participants changed to metal-free clothes and were placed comfortably on the fMEG device. The shielded room was then closed, but participants were constantly monitored from outside through a camera and bidirectional communication was possible via an intercom system. After each stimulation block, participants had the opportunity to take a short break if required. If both stimulation blocks were recorded successfully and participants still felt comfortable on the device, an additional 15 min silent measurement of spontaneous activity was added (which was not included in the current analysis).

### 2.4. Data Analysis

To evaluate fetal brain activity, interfering sources have to be removed from the signal. The main interfering sources in fetal recordings are the maternal and the fetal heart activity. Maternal and fetal heartbeats were detected with the FLORA algorithm (Sippel et al., 2019a) and subsequently removed from the signal, using the FAUNA algorithm (Sippel et al., 2019b). The remaining signal was then used to isolate fetal brain activity. The detailed selection process of channels containing brain activity is described in the supplementary material. In brief, this selection contains the identification of a cluster of channels with the highest amplitudes over one recording block. To ensure that this cluster does not represent artifacts in the data, components containing muscle activity or leftover heart activity are removed beforehand. Regarding the size of the fetal head with respect to the sensor array, a subset of 10 channels is selected during this process. All processing steps were implemented in Matlab R2016b (The MathWorks, Natick, MA).

Once data processing was complete, 122 datasets from 81 different recordings out of the initial number of 202 datasets were available for further analysis. Forty-one recordings retained both datasets (both ssss and sssd rule blocks), 40 recordings were left with only one dataset (17 from the ssss block and 23 from the sssd block). Fifty-six of the datasets had the 500Hz tone as standard tone and 66 the 750Hz tone.

Data from the 10 selected channels per recording were bandpass filtered from 1-10Hz and the root mean square (RMS) over these channels was calculated. The continuous recordings were time-locked to the onset of the four tones sequences and cut into trials from 200ms before the onset of a sequence until 3000ms afterwards (Figure 2). To evaluate an equivalent number of trials per condition, only the standard trials before the deviant trials were used, resulting in N=46 trials per condition. Trials were averaged, resulting in two averages per dataset.

**Figure 2:**
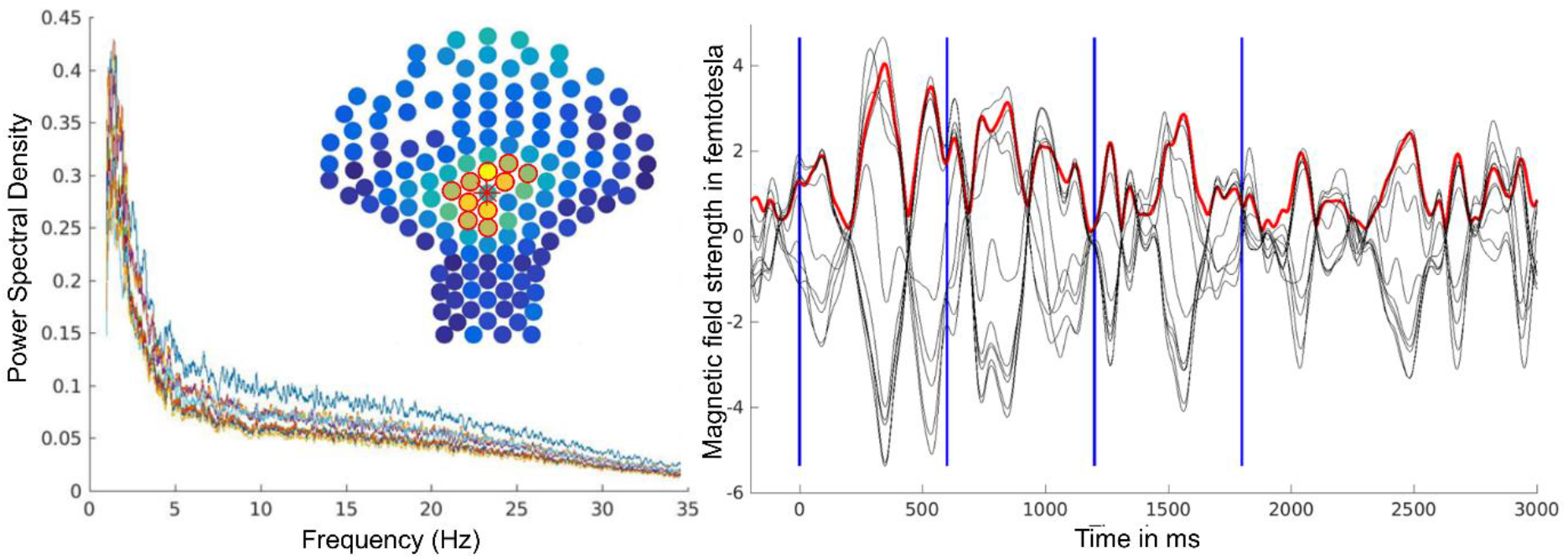
Example of AER towards tone sequence (all trials) of one participant (GA 38 weeks). Left: Cluster of 10 channels within sensor array (marked by red circles) and power spectral density of signal at these channels. Color map represents RMS activity over the time window of one whole recording block. Right: Auditory event-related response (10 channels; black), red line represents RMS, blues lines mark onset of each tone.

#### 2.4.1. Data Normalization

A number of factors, such as the fetal position during the recording, the distance between the fetal head and the sensor array, the fetal head and brain size during different gestational weeks and the number of components removed from the signal during preprocessing, cause a wide range of amplitudes in the trial averages. To overcome this problem we normalized the data. To determine the amount of variability and to optimize the signal-to-noise ratio, we took the trial averages over all 210 trials (learning and testing phase). We then calculated the maximal amplitude of the auditory event-related response (AER) towards the first tone of a sequence for all datasets (Figure 3). Since it follows the long inter-stimulus-interval, this AER towards the first tone is assumed to be the most pronounced response. The amplitude of the AER was determined by the maximum value within 50-350ms following the onset of this first tone of a sequence, which is the usual time window for fetal AERs (e.g. Linder et al., 2014). Since maximal amplitudes span a wide range of values in a non-normally distributed fashion, we normalized data from the two averages per dataset by dividing them by the maximal amplitude of this dataset. The obtained values were then additionally multiplied by 100 to capture the signal in percent signal change from the maximal amplitude of a dataset.

**Figure 3:**
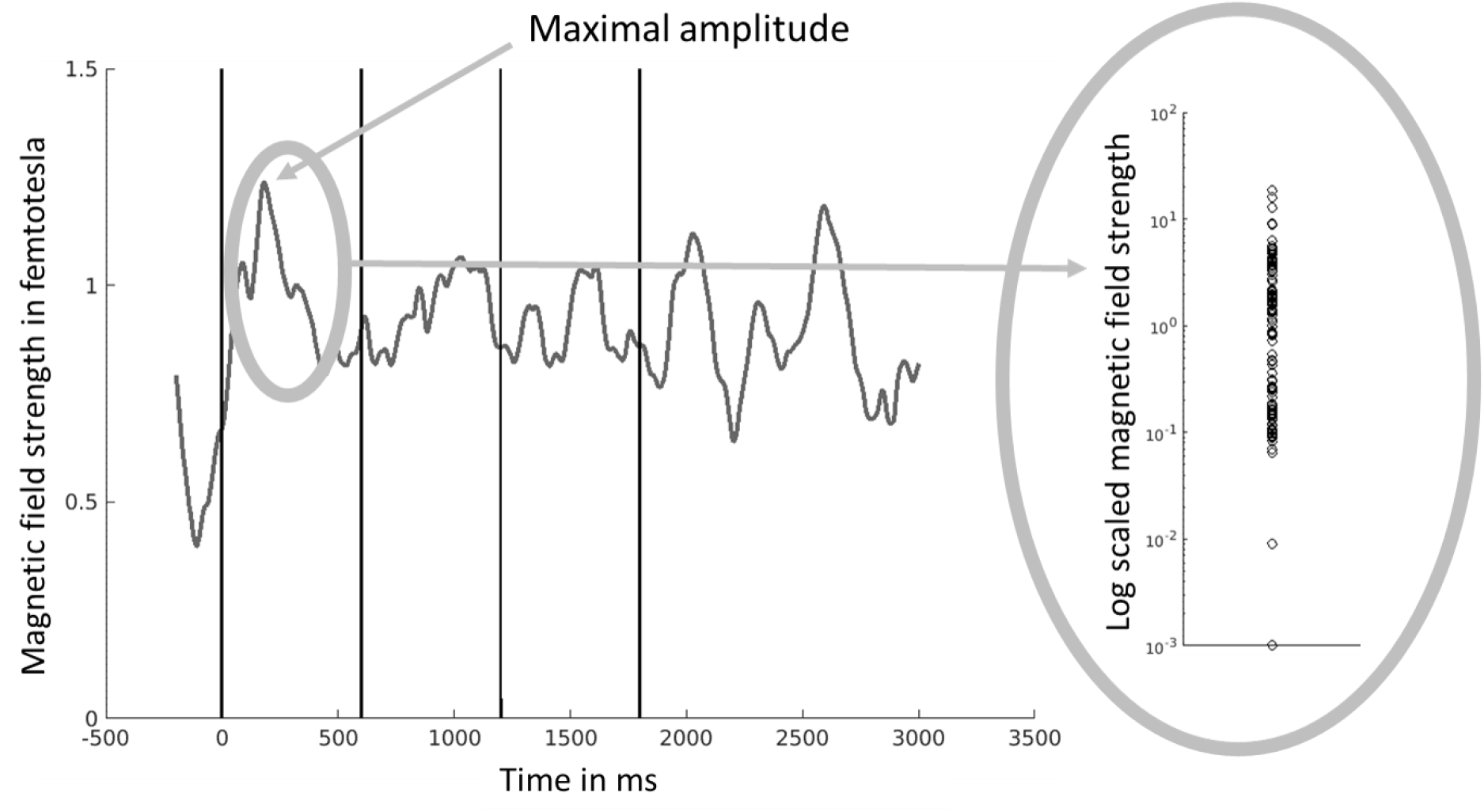
Average event-related response towards all tone sequences (210 trials) over all subjects. Black lines represent onset of each tone. Right graph: Representation of large variability of values of maximal amplitude of first event-related response (50-350ms) for each individual dataset.

To obtain values for further statistical comparisons, two average values per tone were constructed; one for each AER time window (50-350ms after tone onset) and an additional late AER time window 350-650ms after onset of the tone. Other time windows of interest are certainly possible but as there is no prior fetal literature pointing us to a particular time window of interest, we omitted inflating degrees of freedom by constructing additional arbitrary time windows. To detect fetal mismatch responses, AERs towards tone 3 (T3) and tone 4 (T4) of a sequence were compared in both time windows. Response amplitudes towards subsequent tones are usually declining and increase only in the event of a mismatch response (Muenssinger et al., 2013). This approach for detecting fetal mismatch responses enables us to establish a difference value within a sequence, thereby minimizing the impact of variability between datasets (for a summary of the analysis steps see supplementary analysis scheme). Even though restricting our analysis to within-sequence comparisons excludes the possibility to look at late effects after 650ms, the more conservative approach of using within sequence comparisons strengthens the interpretability of our data.

#### 2.4.2. Analysis of Heart Rate Variability (HRV)

Fetal HRV was calculated from the R peaks of the fetal magnetocardiogram detected during data preprocessing. For N=7 datasets the R peak detection was not of sufficient quality for HRV analysis, resulting in N=115 datasets including HRV. As in Moser et al. (2020), we computed the standard deviation of normal to normal R-R intervals (SDNN) as the HRV parameter for each dataset, using in-house algorithms as described in Mat Husin et al. (2020). In accordance with the analysis presented in this paper, we created high HRV and low HRV extreme groups, taking the highest third and lowest third of datasets into account and omitting the datasets with a medium HRV value.

### 2.5. Statistical Analysis

All statistical analyses were performed in R (R Core Team, 2019). As a first analysis step, a two-sided paired t-test was used to compare responses to T3 and T4 in each condition. In a second step, differences between T4 and T3 (T4-T3) were compared between conditions. These comparisons were carried out with a mixed model (lme4; Bates et al., 2015). This method was selected since data in all four experimental conditions were not available in all recordings and mixed models are a recommended method for dealing with missing values. Data normal distribution was given as a result of the previous data normalization step.

Recording number was used as a random effect and the role of the condition in either the local or the global regularity as a fixed effect (each with two levels). Significances of the mixed models were tested with a type II Analysis of Variance with Satterthwaite’s method to estimate effective degrees of freedom (lmerTest; Kuznetsova et al., 2017). A Quantile-Quantile Diagram was used to ensure normal distribution of the model residuals. For post-hoc testing of the differences between individual conditions estimated marginal means were calculated. Each test was performed for both, the early and the late AER window.

For analyzing the impact of gestational age on responses towards local and global rule violations, participants were separated into an early (weeks 25-34) and a late gestation (weeks 35-40) group. To obtain similar groups sizes the selection of time ranges was data driven. First, GA group was added as an additional fixed effect and interaction term (global rule* GA group) to the model described above. Conditions were later compared in each of the groups as described above (N=68 datasets early GA and N=54 datasets late GA with local rule or global rule as a fixed effect and performing post-hoc tests on the conditions). To obtain a more fine-grained picture of the effect of GA, Pearson correlations between GA and the difference T4-T3 were calculated for each condition. Additionally, the differences between T4-T3 in the sssD and ssss condition (in the block with the ssss rule) and the differences between T4-T3 in the sssS and sssd condition (in the block with the sssd rule) were correlated with GA. These correlations provide insight into the development of local vs. global rule learning over gestation.

For the exploration of the impact of behavioral state on global rule learning, datasets from the late gestation group only were taken into consideration. Since this group contains 54 datasets, we used N=18 datasets with a high SDNN (upper third) for the high HRV group and N=18 datasets with a low SDNN (lower third) for the low HRV group. Conditions were compared in each of the groups as described above (using global rule as fixed effect and doing post-hoc tests on the conditions). In all tests, the significance level was set to α=0.05 and was not corrected for multiple comparisons. Due to the complexity of fetal data and the lack of similar previous studies we were unable to formulate precise hypotheses. We would therefore emphasize that this is an explorative rather than a confirmatory analysis. The usage of uncorrected p-values in this exploratory context is a matter of some debate (Rubin, 2017).

## 3. Results

### 3.1. Hierarchical Rule Learning Over Gestation

In the early AER time window (50-350ms), participants did not display any differences between responses towards T3 and T4 or any effect of local or global rule violations when comparing these T4-T3 differences between conditions. All the following results therefore relate to the late AER time window (350-650ms). First, we compared responses towards T3 and T4 for each individual condition. Only one condition (sssD) revealed a significantly different response (t(57) = 2.51, p = 0.015), showing an increase of event-related activity from T3 to T4 (supplementary Table 2). Second, we compared the differences T4-T3 between the four stimulation conditions. Participants showed a significantly higher difference for those conditions that violate the global rule (F(1,173.36) = 4.56, p = 0.034). There was no effect of local rule (supplementary Table 4). Pairwise comparison of individual conditions revealed that only the sssd and sssD conditions differed significantly (t(203) = 2.31, p = 0.022; Figure 4a; supplementary Table 6). When the GA groups (early gestation and late gestation) were included in the model for global rule violations, the main effect for the global rule remained significant (F(1,172.44) = 4.58, p = 0.034) and an additional marginally significant effect for the GA group (F(1,73.39) = 3.92, p = 0.051), but no interaction (F(1,172.44) = 2.25, p = 0.14) was observed.

**Figure 4:**
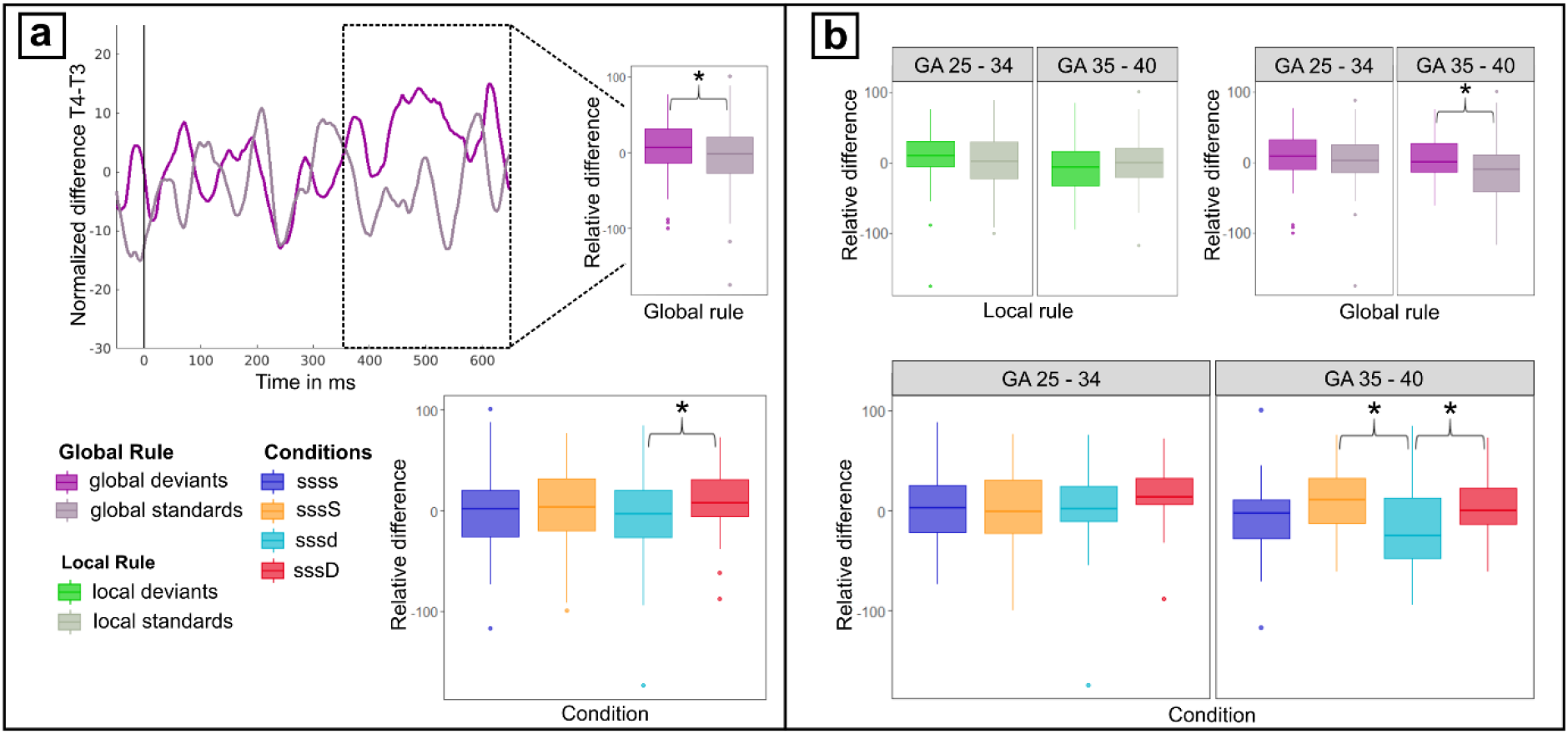
Differences tone 4 - tone 3 in time window between 350-650ms for each stimulation condition. a) time course of differences for global standards and global deviants with average responses in time window of interest; b) split of data depicted in a) into two gestational age groups: early gestation (GA 25-34 weeks) and late gestation (35-40 weeks). * depict significant differences p<0.05.

This led us to the follow-up analysis, which involved testing each of the GA groups individually. The previous finding, which showed a significant response increase from T3 to T4 in the sssD condition, was also present in the early GA group (t(29) = 2.8, p = 0.009) but was, interestingly not observed in the late GA group (t(27) = 0.69, p = 0.494). The late GA group showed a significant decrease from T3 to T4 in the sssd condition only (t(25) = -2.14, p = 0.042; supplementary Table 3). As already carried out with the whole group, we compared the differences T4-T3 between the four conditions. The difference between globally standard and globally deviating conditions was no longer significant in the early GA group (F(1,134) = 0.35, p = 0.556) and was more pronounced in the late GA group (F(1,74.43) = 6.83, p = 0.011; Figure 4b; supplementary Table 5). The post-hoc test for individual conditions showed that in the late GA group conditions sssS-sssd and sssD-sssd differed significantly (t(72.8) = 2.39, p = 0.012; t(84.2) = 2.21, p = 0.03; supplementary Table 7).

With regard to the correlation between GA and the difference T4-T3 for each individual condition we detected a significant negative correlation for the sssd condition (r(62)=-0.319, p = 0.01; Figure 5) but no significant correlations for any of the other conditions. Upon assessment of the correlation between the effect of the global rule and GA in each block (ssss rule and sssd rule), we observed a marginally significant correlation for the difference T4-T3 between sssS-sssd with GA (r(62) = 0.216, p = 0.087) but nothing for sssD-ssss (r(56) = 0.048, p = 0.719; Figure 5).

**Figure 5:**
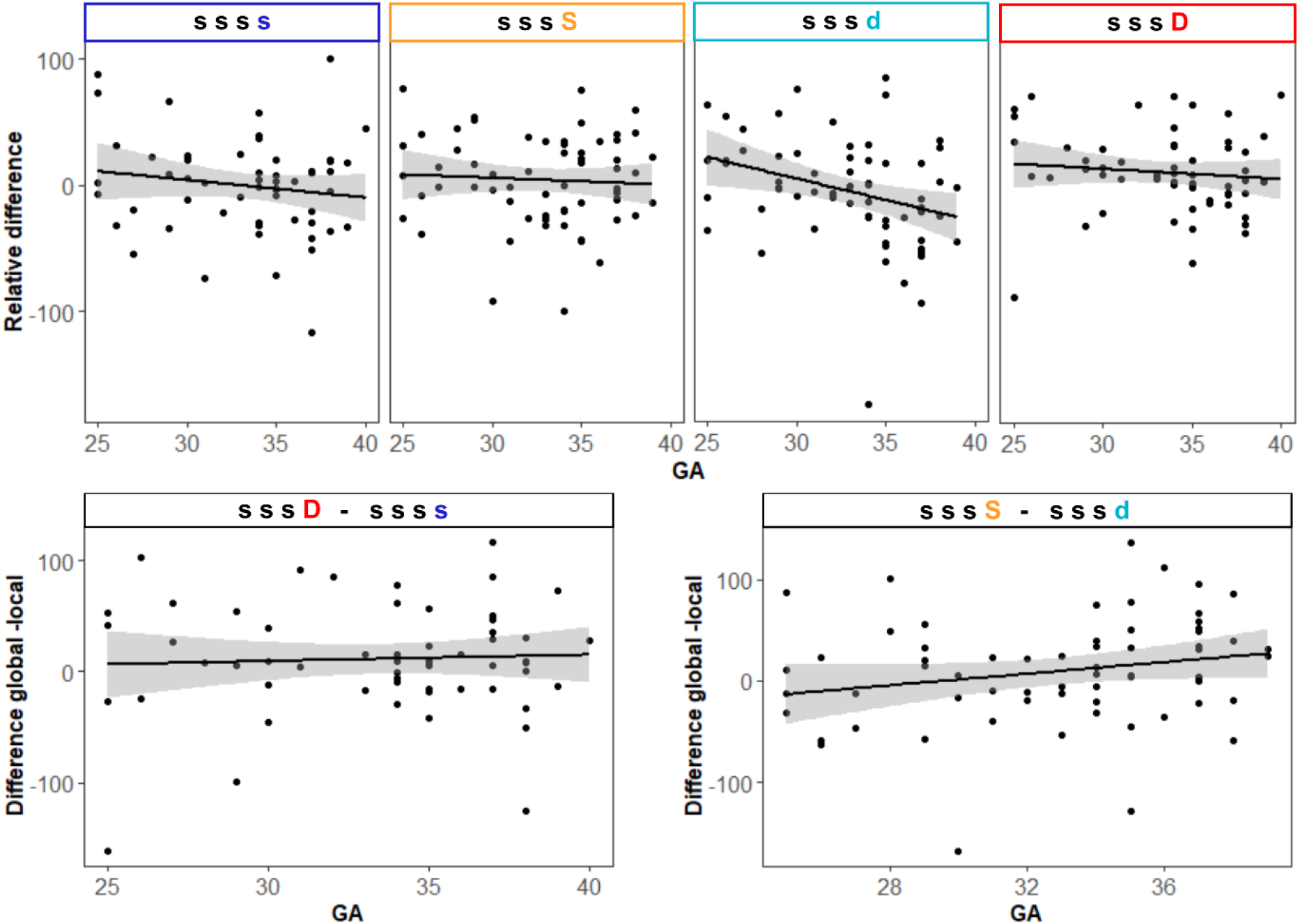
Top row: Correlation of difference tone 4 - tone 3 and gestational age. Significant correlation was detected for sssd condition only (p<0.05). Bottom row: Correlation between difference tone 4 – tone 3 for left: sssD-ssss and right: sssd-sssS (marginally significant correlation, p<0.1).

### 3.2. Impact of Fetal Behavioral State

Fetal HRV (used as proxy for behavioral state) in the late GA group (N=54) ranges from an SDNN of 4.14 to 36.04 with a median of 9.48. The SDNN in the high HRV group (N=18) ranges from 10.79-36.04 with a median of 15.18 and in the low HRV group (N=18) from 4.14-8.3 with a median of 6.14. The evaluation of the two different groups revealed that fetuses in the high HRV group showed a highly significant effect of the global rule (F(1,34) = 9.33, p = 0.004) while fetuses in the low HRV group showed no significant effect of the global rule (F(1,19.28) = 0.69, p = 0.417; Figure 6). The post-hoc test for individual conditions in the high HRV group showed that conditions sssS-sssd and sssD-sssd differed significantly (t(32) = 2.74, p = 0.01; t(32) = 2.49, p = 0.018) and that sssS-ssss differed marginally significantly (t(32) = 1.77, p = 0.087).

**Figure 6:**
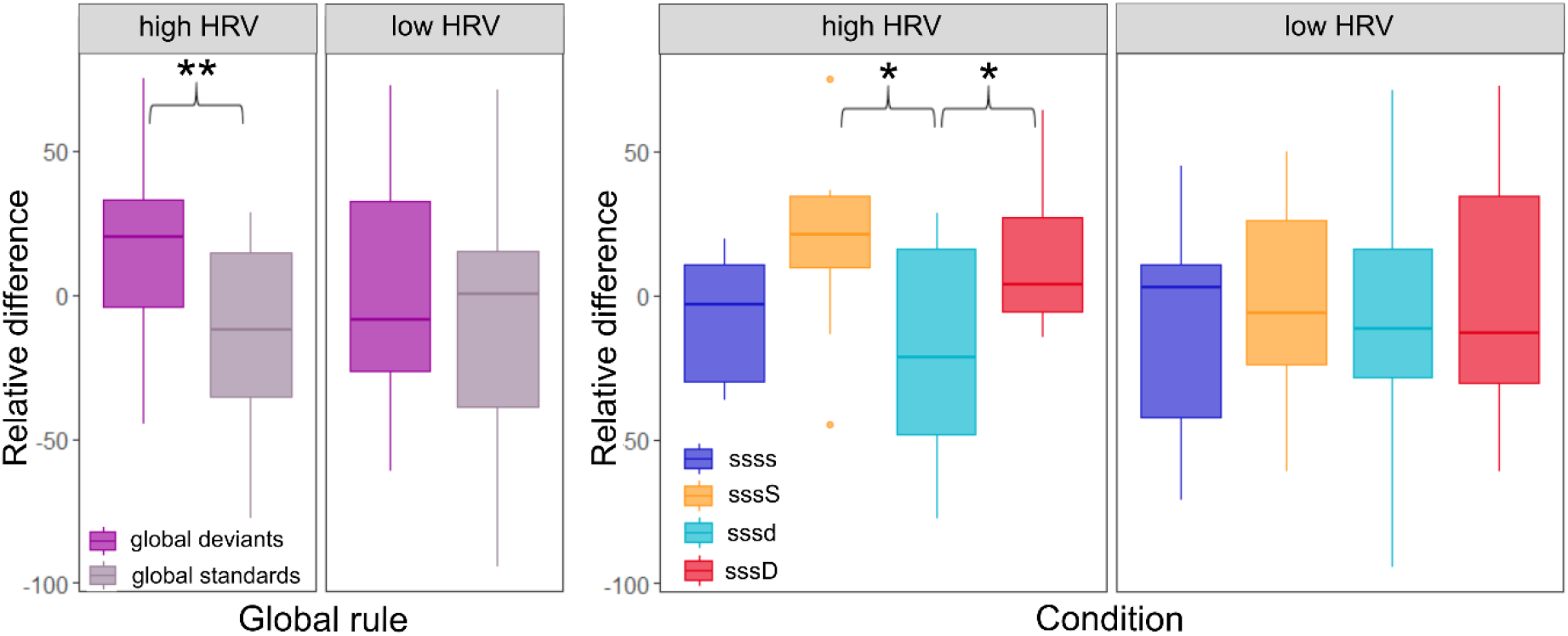
Difference tone 4 - tone 3 in time window between 350-650ms for each stimulation condition in fetuses in late gestation (GA 35-40) with either high or low HRV values. Left: effect of global rule, right: individual conditions. * depicts significant difference, ** highly significant.

## 4. Discussion

The main insight of the present study is that fetuses in the third trimester of gestation show signs of hierarchical rule learning. This was demonstrated by event-related responses in a time-window between 350-650ms after stimulus onset. The difference in activation from a standard third tone to a fourth tone, violating the second-order regularity, was significantly higher than the difference to a fourth tone, following the second-order regularity, as shown by a main effect for second-order regularity in the comparison of these differences. Separating the early and late gestation half of the sample revealed that this effect was mainly driven by the fetuses in later gestational weeks. The effect for the second-order rule violation was more pronounced in the group of older fetuses than in the whole sample and the younger fetuses did not show any rule-learning effect. Within the older group, a comparison between fetuses in different behavioral states showed that the hierarchical rule-learning effect was mainly shown by the fetuses in a more active behavioral state, expressed by a high HRV.

### 4.1. Fetal Hierarchical Rule-learning Ability

The investigation of the two time windows after tone onset (50-350ms and 350-650ms) revealed that in the early window – previously defined as the window of the auditory event-related response – participants did not differentiate between stimulation conditions at all. This shows that at the early AER stage, it is too soon for a fetal mismatch response. The differences observed in the later time window cannot clearly be assigned to either the mismatch response or to the late discriminative negativity component, as they do not fit into the time windows found in infant studies (200-400ms for mismatch, and after 680ms for the late component (Dehaene-Lambertz and Dehaene, 1994; Háden et al., 2016). However, since there are no known clear component structures in fetuses, we had no clear prior expectations on the time windows of interest. Unfortunately, restricting the analysis on the differences between T4-T3 did not permit an investigation of time windows later than 650ms after the onset of T4, which could probably have revealed additional late effects. However, this restriction was important to ensure an optimal signal-to-noise ratio for the effects investigated.

The direct comparison between T3 and T4 was significant for the local and global deviant (sssD) only. We had anticipated that this would be the strongest response since it is the rarest deviant within the paradigm and since the amplitude of a mismatch response increases with decreasing stimulus probability (Baldeweg et al., 2004). When looking at the early and late gestation group separately we found that this direct comparison in the sssD condition was only significant in the early gestation group. The most likely explanation for this curious finding is that the timing of the mismatch response changes over the course of gestation and the response is not captured well by the 350-650ms time window in the late gestation group. Our analysis approach with one early and one late time window does unfortunately not allow for a more detailed investigations of such potential changes in timing. Lack of responses for other deviants could relate to the fact that we used RMS values over rather large time windows. Previous fMEG studies, using pre-defined time windows to investigate peaks of mismatch responses, also reported responses in only 60-66% of their sample (Draganova et al., 2005; 2007). Since studies in preterm infants reported responses towards first-order deviants as early as between week 28 and 32 (Mahmoudzadeh et al., 2017), first-order rule learning should be generally present.

### 4.2. Developmental Trend of Second-order Rule Learning

When dividing the sample into an early and a late gestation group, we ascertained that mismatch responses towards the local and global deviant were present already at an early stage of the last trimester of pregnancy, but that second-order rule learning could be detected only in the late gestation group. As observed in our correlation analysis, the response towards the sssD sequence does not change over gestation. The response towards the other two types of deviants (sssd and sssS), however, reverses its pattern from high for sssd in the early weeks to a comparably elevated response towards sssS in the later weeks. This shows a trend towards an increase in second-order rule-learning abilities over the course of gestation. However, this trend was mainly driven by a decrease in the response towards the sssd deviant.

During the development of the human brain in late gestation, multiple processes could facilitate the ability of second-order rule learning. Thalamo-cortical connections are more mature and show increased myelination, and cortico-cortical connections grow increasingly (Kostović et al., 2019). This could lead away from the very modular structure of the young brain toward a more integrated one. Weeks 35-37 of gestation are particularly dominated by the growth of interhemispheric connections (Kostović and Judaš, 2010). In addition, the connections between the posterior cingulate cortex and other brain regions show a developmental trend over the last trimester of gestation. This constitutes a sign for a decrease in modularity and an increase in information processing competencies as the posterior cingulate cortex is a connective hub for a small-world network organization (Thomason et al., 2014). Generally speaking, the brain structure becomes more globally connected during development. However, even small children show small-world network organization, meaning that networks allow both local and distributed processing, which is a sign for efficient network communication (Fair et al., 2009). In fetuses older than 35 weeks of gestation, connections between regions known to belong to the DMN can already be observed (Thomason et al., 2015). This network is important for temporal awareness and integration of perceptual consciousness (Kent et al., 2019). This increase in connectivity can be seen as a driving factor to make conscious perception possible, e.g., when referring to the global workspace theory of consciousness (Baars, 1997; Dehaene et al., 1998). Work with adult volunteers suggests that conscious awareness is mediated by two distinct networks, one for external awareness (containing lateral fronto-parietal areas) and one for internal awareness (largely overlapping with the DMN; Vanhaudenhuyse et al., 2011). These network showed to be disconnected in patients in an unresponsive state and start to reconnect once patients recover to a minimally conscious state (Demertzi et al., 2013).

On the whole, our results show that fetuses older than week 35 GA were able to make predictions at a temporal scale that matched the underlying structure of the paradigm, which can be seen as a sign of consciousness (Marchi and Hohwy, 2020). We cannot, of course, establish the beginning of consciousness at the 35^th^ week of gestation on the basis of our results, as our division of the fetuses into two groups was driven by the data availability. What our results do show, however, is that there is a linear trend with gestation and we will certainly have to consider inter-individual differences in the future.

One interesting aspect of the results is the decrease of neural responses towards local rule violations over gestation. This could be indicative of an increase in the fetal ability to represent the second-order rule. In this case, the memory for the sssd sequence as the second-order rule would impact the predictions formed by the fetus and lead to a decrease in reactivity towards the fourth tone in accordance with this rule. However, when contemplating the results, we also have to bear in mind that only the standard sequences before each deviant sequence were evaluated, meaning that the sequence had already been repeated multiple times. Responses may well look different once all local deviants, especially those in the learning phase, are evaluated.

### 4.3. Impact of Activity State

The analysis on the impact of fetal behavioral state on second-order rule learning was performed on the basis of results from the previous study with newborns (Moser et al., 2020). As expected, effects resembled those obtained in Moser et al. (2020), showing that the effect for the global rule in fetuses in late gestation was clearly present in those in a more active (high HRV) state, while it was not detectable in those fetuses in a more quiet (low HRV) state. Replication of this effect shows that the impact of behavioral state on cognitive processes accounts for both newborns in their first week of life and fetuses in the last weeks of gestation, regardless of whether they are in- or outside the womb. The similarity of results in fetuses and newborns emphasizes the role of behavioral state for learning in early life. Furthermore, results are interesting in the light of discussions on how much sedative placental hormones in the fetal environment suppresses fetal wakefulness (Lagercrantz and Changeux, 2009; Padilla and Lagercrantz, 2020). The difference in responses shown by fetuses in different behavioral states challenges the assumption that fetuses are mostly asleep and that the fetal environment suppresses conscious processing of stimulation from outside the womb. The two dimensions – maturational stage (gestational age) and activity state – impacting our findings highlight a particular challenge of this work compared to work with adult populations. Unlike in fetuses, when studying adult populations, there is a clear assumption about the ability to consciously process stimuli before entering an altered state of consciousness for example during anesthesia (e. g. Sanders et al., 2012) or caused by brain damage (Gosseries et al., 2014). Nevertheless, all these cases face the challenge of quantifying conscious processing without a possibility for communication.

### 4.4. Limitations and Outlook

Even though, this study provides empirical evidence for conscious processing of stimuli from outside the womb, it is important to mention that the paradigm implemented in this study only covers one of the many aspects of consciousness. As laid out in the introduction, this approach focuses on a more primary form of consciousness and is not related to concepts like self-consciousness. Even though we gained insights into the complex cognitive processes that start to develop before birth, we are still far from drawing conclusions about the qualia of fetal experiences.

One major limitation of the presented study is the data quality of fMEG recordings. The poor signal-to-noise ratio together with the limited amount of recordings where brain activity can be extracted poses challenges for data analysis. The present analysis focused on methods to minimize processing-related variance and was therefore limited to certain aspects of the fetal responses. Further methodological developments could improve data processing and enable us to consider additional aspects of the data such as, for example, late fetal responses. Furthermore, prior knowledge of fetal mismatch responses is limited and the time windows of interest are influenced by parameters such as the gestational age and the behavioral state. The simplistic approach of studying one early and one late time window was chosen. A more fine-grained view on the temporal dynamics of responses could help to develop hypotheses for further mismatch studies. Additionally, other parameters such as power spectral density or signal complexity, neither of which are dependent on an exact time window could be observed in further analyses (Ruffini, 2017; Tononi and Edelman, 1998).

It should moreover be mentioned that based on the nature of fMEG recordings it is not possible to investigate fetuses independent of their mothers. Mothers were also exposed to the sound stimulation and therefore may have reacted to the tones (e.g. by moving or tensing muscles). However, in addition to clear instructions to ignore the tones, our pre-processing steps ensured that maternal muscle artifacts were removed and datasets with numerous movement artifacts were removed from the analysis entirely. Therefore, we can attribute the reactions found with a high degree of certainty to the fetus and not to the mother.

Our initial dataset contained multiple longitudinal recordings over four time-points during pregnancy. However, the aforementioned challenges in data processing rendered it impossible to investigate differences in individual trajectories. Acquisition of even more time-points during gestation would help to compensate for data lost during later processing steps. In general, availability of larger datasets would make it easier to compensate for the limited data quality. New developments in the field of optically pumped magnetometers could increase the availability of fetal data. These sensors operate without the intensive cooling required for MEG sensors and therefore do not require the complex infrastructure of fMEG devices, which would allow to use them in a greater number of recording facilities (Boto et al., 2017). In addition, hierarchical rule learning could be tested in preterm infants in equivalent age groups. As auditory oddball paradigms are well established in studies with preterm infants (e.g. Mahmoudzadeh et al., 2017) the local-global paradigm could be easily transferred to this group. Even though fetuses and preterms at the same gestational week can not be treated equivalently, high quality recordings in preterms can contribute to the understanding of the development of conscious processing.

## 5. Conclusions

In a nutshell, the current study shows that fetuses in the final weeks of gestation are able to learn second-order regularities. This ability is considered to be a sign for conscious processing and therefore raises the point that consciousness probably does not start with birth. The brain structural basis for this ability is already available with the establishment of thalamo-cortical connections around week 25 of gestation, although the current study shows that it gradually develops further during the last trimester. This development could be related to a general increase of brain connectivity to allow a more global processing.

## 6. Acknowledgements

We wish to thank Janina Einsele for her support with data acquisition. This work was funded by the FET Open Luminous project (H2020 FETOPEN-2014-2015-RIA under agreement No. 686764) as part of the European Union’s Horizon 2020 research and training program 2014–2018 and the German Federal Ministry of Education and Research (BMBF) to the German Center for Diabetes Research (DZD e.V. 01GI0925). The authors thank the International Max Planck Research School for the Mechanisms of Mental Function and Dysfunction (IMPRS-MMFD) for supporting Lorenzo Semeia.

## 7. Author Contributions

*Julia Moser:* Investigation, conceptualization, methodology, formal analysis and writing – original draft. *Franziska Schleger:* Conceptualization, methodology and writing – review & editing. *Magdalene Weiss:* Investigation and data curation, project administration. *Katrin Sippel:* Methodology and software. *Lorenzo Semeia:* Formal analysis and writing – review & editing. *Hubert Preissl:* Supervision, conceptualization and writing – review & editing. All authors read and approved the final version of the manuscript.

