## Supplementary Material for "Magnetoencephalographic Signatures of Conscious Processing Before Birth"

**This Supplementary information includes:**

Tables S1 to S7
Supplementary information text: Fetal brain detection
Supplementary figure: Data analysis scheme

**Supplementary Tables**

**Table 1.** Number and distribution of participants

| Participants | Longitudinal participation | Recordings | Datasets |
|---|---|---|---|
| N=60 | | | |
| **Included: N=56** | 1 x N=34<br>2 x N=7<br>3 x N=3<br>4 x N=12 | N=105<br>GA 25-28: N=23<br>GA 29-32: N=24<br>GA 33-36: N=35<br>GA 37-40: N=23 | N=202 |
| **Included in brain analysis: N=43** | 1 x N=24<br>2 x N=5<br>3 x N=9<br>4 x N=5 | N=81<br>GA 25-28: N=14<br>GA 29-32: N=18<br>GA 33-36: N=28<br>GA 37-40: N=21 | N=122<br>ssss & sssd block: N=40<br>ssss only: N=17<br>sssd only: N=23 |

**Table 2.** Differences from tone 3 to tone 4 (T4-T3)

| Condition | mean difference | df | t-value | p-value |
|---|---|---|---|---|
| **ssss** | -0.63 | 57 | -0.13 | 0.901 |
| **sssD** | 10.71 | 57 | **2.51** | **0.015** |
| **sssd** | -4.99 | 63 | -0.92 | 0.359 |
| **sssS** | 4.24 | 63 | 0.95 | 0.345 |

**Table 3.** Differences from tone 3 to tone 4 (T4-T3) for different gestational age groups

| Condition | mean difference | df | t-value | p-value |
|---|---|---|---|---|
| | | **Early gestation (GA 25-34)** | | |
| **ssss** | 5.18 | 29 | 5.18 | 0.458 |
| **sssD** | 16.9 | 29 | **2.8** | **0.009** |
| **sssd** | 3.99 | 37 | 3.99 | 0.557 |
| **sssS** | 1.61 | 37 | 1.61 | 0.792 |
| | | **Late gestation (GA 35-40)** | | |
| **ssss** | -6.87 | 27 | -0.93 | 0.359 |
| **sssD** | 4.08 | 27 | 0.69 | 0.494 |
| **sssd** | -18.12 | 25 | **-2.14** | **0.042** |
| **sssS** | 8.1 | 25 | 1.23 | 0.231 |

**Table 4.** Effects of local and global rule and gestational age on differences T4-T3

| Rule/Effect | Sum of squares | df | F-value | p-value |
|---|---|---|---|---|
| **Global rule** | 6392.9 | 1, 173.36 | **4.55** | **0.034** |
| **Local rule** | 18.363 | 1, 173.88 | 0.01 | 0.91 |
| **GA group** | 5472.4 | 1, 73.39 | 3.92 | 0.051 |
| **Global rule * GA group** | 3148.0 | 1, 172.44 | 2.25 | 0.135 |

**Table 5.** Effect of local and global rule on differences T4-T3 in the two gestational age groups

| Rule | Sum of squares | df | F-value | p-value |
|---|---|---|---|---|
| | | **Early gestation (GA 25-34)** | | |
| **Global rule** | 500.21 | 1, 134 | 0.35 | 0.556 |
| **Local rule** | 1435.5 | 1, 134 | 1.01 | 0.318 |
| | | **Late gestation (GA 35-40)** | | |
| **Global rule** | 9040.7 | 1, 74.432 | 6.84 | 0.011 |
| **Local rule** | 1301.5 | 1, 75.016 | 0.91 | 0.342 |

**Table 6.** Pairwise comparison with estimated marginal means for differences T4-T3 in each condition

| Contrast | estimate | df | t-value | p-value |
|---|---|---|---|---|
| **sssD-ssss** | 11.34 | 172 | 1.63 | 0.105 |
| **sssD-sssd** | 15.72 | 203 | **2.31** | **0.022** |
| **sssD-sssS** | 6.49 | 203 | 0.95 | 0.342 |
| **sssS-sssd** | 9.23 | 172 | 1.39 | 0.166 |
| **sssS-ssss** | 4.85 | 203 | 0.71 | 0.477 |
| **sssd-ssss** | -4.38 | 203 | -0.64 | 0.521 |

**Table 7.** Pairwise comparison with estimated marginal means for differences T4-T3 in each condition for different gestational age groups

| Contrast | estimate | df | t-value | p-value |
|---|---|---|---|---|
| | | **Early gestation (GA 25-34)** | | |
| **sssD-ssss** | 11.71 | 132 | 1.2 | 0.232 |
| **sssD-sssd** | 12.91 | 132 | 1.4 | 0.164 |
| **sssD-sssS** | 15.29 | 132 | 1.66 | 0.1 |
| **sssS-sssd** | -2.38 | 132 | -0.28 | 0.784 |
| **sssS-ssss** | -3.58 | 132 | -0.39 | 0.699 |
| **sssd-ssss** | -1.20 | 132 | -0.13 | 0.897 |
| | | **Late gestation (GA 35-40)** | | |
| **sssD-ssss** | 10.95 | 72.8 | 1.12 | 0.266 |
| **sssD-sssd** | 22.07 | 84.2 | **2.21** | **0.03** |
| **sssD-sssS** | -4.14 | 84.2 | -0.42 | 0.679 |
| **sssS-sssd** | 26.21 | 72.8 | **2.59** | **0.012** |
| **sssS-ssss** | 15.09 | 84.2 | 1.51 | 0.134 |
| **sssd-ssss** | -11.12 | 84.2 | 1.12 | 0.268 |

**Supplementary Information Text**

**Fetal Brain Detection**

For the evaluation of fetal brain activity, it is necessary to detect clusters of brain activity within the SARA sensor array. Those clusters usually span around 10 sensors and are located near the position of the fetal head. In theory, those sensors should be the ones with the highest activity after removal of the maternal and fetal heart activity. Yet, detection of fetal brain activity can be limited by additional interfering sources left in the signal. One possible source of interference is maternal tight muscle activity as the tights of the mother are directly next to the sensors in the lower part of the sensor array. If a participants' position on the SARA system is not optimal or gets uncomfortable during the recording time, this is often accompanied by rising tension in tight muscles. Another possible source is leftover of fetal or maternal heart activity. As the heart activity is up to 10000 times stronger than the fetal brain activity, it is possible that there are leftovers after FAUNA removal.
The following part describes the brain detection algorithm:

**Sensor check.** As a first step, the signal is checked for disturbing sensors. For the sensor check, the variance of each sensor and the correlation coefficients between each sensor and its neighbor sensors are calculated. Neighbor sensors are defined as sensors with less than 5 cm distance. To be included into further evaluation, the variance of the signal within each sensor has to be below 10% calculated on the normalized data (normdata = (data - min(data)) / (max(data) - min(data)) and the correlation r-value between a sensor and its neighboring sensors has to be above 0.1 and significant after Bonferroni correction of the significance level (0.05 divided by the number of sensors).

**PCA and removal of interfering components.** To account for possible interfering sources, the magnetic signal is split into its principal components and checked for components that belong to interfering sources. Since we try to identify these components based on their spectral power density (PSD) characteristics, a fast Fourier transform on each component is calculated as a first step.

**Identification of muscle activity.** While brain activity usually shows a 1/f signature, muscular activity in a band-pass filtered power spectrum from 1 to 35 Hz appears as a very spiky shape that can be approximated by an inverse U-shape, starting very low in the lower frequency bands and peaking between 15-20 Hz.

To identify muscular components, a second degree polynomial curve (f(x) = ax^2*bx+c) is fitted to the power spectral density from 1-35 Hz. An inverse shape of this polynomial (function with a negative a) is an indication for muscle activity and the respective component is classified as such.

**Identification of leftover heart activity** For the identification of leftover heart activity, the power spectral density from 1-7 Hz is correlated with three different functions:

- a logarithmic function: *f_brain(xf) = (-log(xf)+2)*0.5,* describing the usual 1/f signature of brain activity, where *xf* is the x-axis of the frequency in Hz.
- a logarithmic cosine function (*f_fmcg*) using the main frequency of the fetal heat signal (RR_f) as it is calculated by the FLORA algorithm: *f_fmcg(xf) = ((cos(2*pi*RR_f* xf)*0.5)+1)*(f_brain).*

- a logarithmic cosine function (*f_mmcg*) calculated the same way as *f_fmcg* using the main frequency of the maternal heat signal (*RR_m*) instead of *RR_f*.

All three functions are by calculation already compressed and transferred to the positive area of the y-axis. If the correlation of a principal component with one of the logarithmic cosine functions is higher than its correlation with *f_brain*, the component is classified as leftover heart activity.

All component showing either muscular or leftover heart-activity are multiplied by zero and therefore removed from the dataset during the process of inverse PCA. After removal of all disturbing components, the cleaned signal should only contain fetal brain activity.

**Brain cluster detection.** As the fetal brain is localized around the fetal head and not distributed over the whole sensor array, we selected a cluster of 10 sensors with the highest activity. Therefore, first the root mean square (RMS) activity of all sensors is calculated and sorted. The RMS is used to account for the dipolar structure of magnetic signals. Then, starting with the sensor with the highest RMS activity all sensors are inspected in descending order. The highest sensor marks the center of the first possible cluster. If the next sensor is within 10cm of this clusters center, the sensor is added to that cluster and the cluster's center recalculated. If the sensor is not fitting to the cluster a new cluster is added. The next sensor is then compared to all possible cluster centers. As soon as one cluster contains 10 sensors the process is ended and this cluster is selected as brain activity.

**Visual examination.** The last processing step contains a visual inspection, in which the rater determines whether the power spectrum of the selected channels reflects approximately a logarithmic function and whether the location of the 10 selected channels fits to the location determined by ultrasound. In case the power spectrum is flat or contained peaks from other sources of noise, datasets are not considered for evaluation. Same applies for a large divergence between the position of the fetal head and the localized cluster. A flat PSD can result from recordings containing weak brain activity in combination with numerous disturbing sources.

**Fig. S1.** Data Analysis Scheme

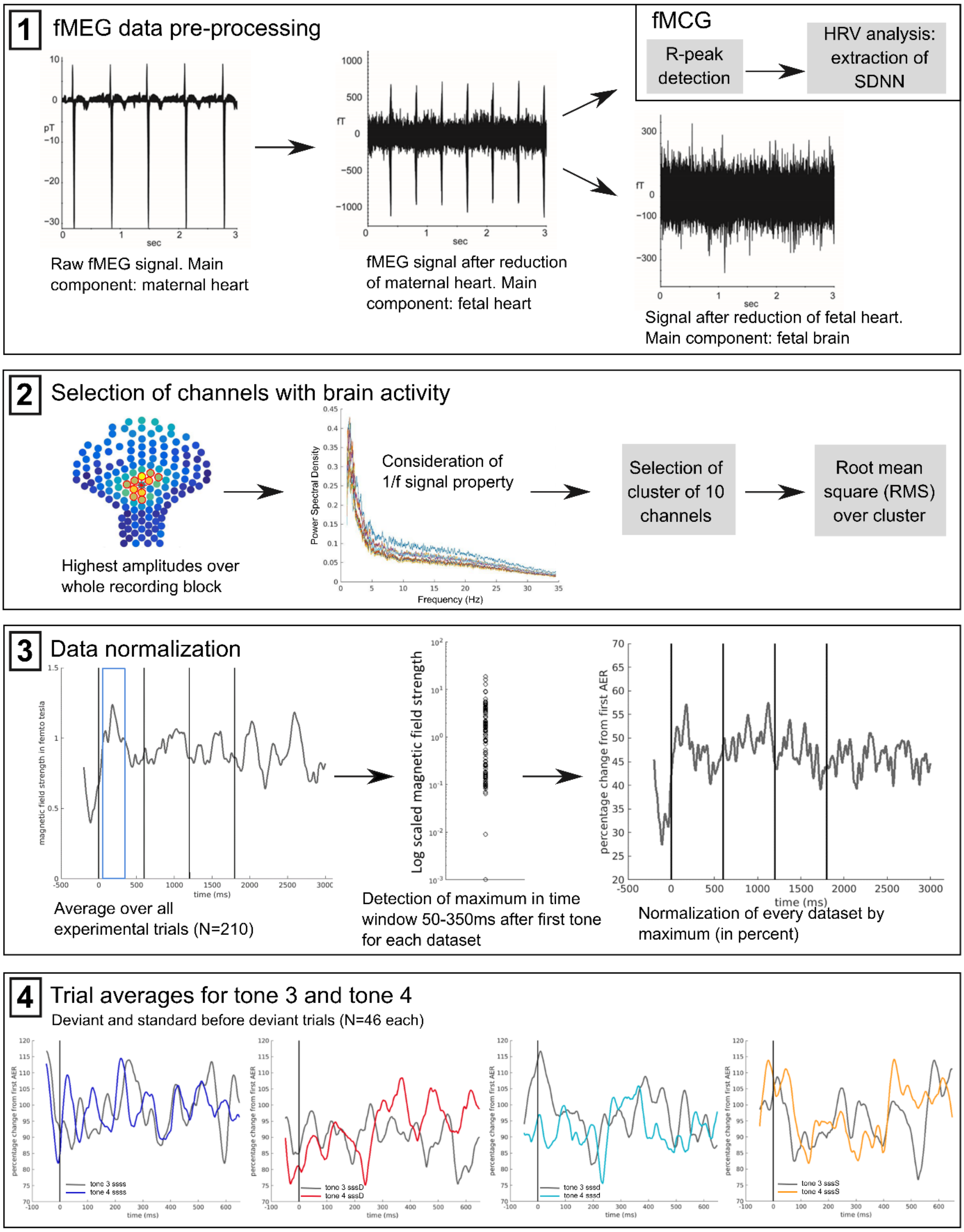
